# Structural mechanism of angiogenin activation by the ribosome

**DOI:** 10.1101/2023.12.11.570495

**Authors:** Anna B. Loveland, Cha San Koh, Robin Ganesan, Allan Jacobson, Andrei A. Korostelev

**Affiliations:** RNA Therapeutics Institute, UMass Chan Medical School, Worcester, MA, USA; Department of Microbiology and Physiological Systems, UMass Chan Medical School, Worcester, MA, USA

## Abstract

Angiogenin, an RNase A-family protein, promotes angiogenesis and has been implicated in cancer, neurodegenerative diseases, and epigenetic inheritance. Upon activation during cellular stress, angiogenin cleaves tRNAs at the anticodon loop, producing nicked tRNA (also known as tRNA halves, tiRNAs or tsRNAs) and resulting in translation repression. The catalytic activity of isolated angiogenin, however, is very low, and the mechanisms of the enzyme activation and tRNA specificity have remained a puzzle. Here, we uncover these mechanisms using biochemical assays and cryogenic electron microscopy. Our work reveals that the cytosolic ribosome is the long-sought activator of angiogenin. A 2.8-Å resolution cryo-EM structure features angiogenin bound in the A site of the 80S ribosome. The C-terminal tail of angiogenin is rearranged by interactions with the ribosome to activate the RNase catalytic center, making the enzyme several orders of magnitude more efficient in tRNA cleavage. Additional 80S•angiogenin structures capture how the tRNA substrate is directed by the ribosome next to angiogenin’s active site, demonstrating that the ribosome acts as the specificity factor. Our findings therefore suggest that angiogenin is activated by ribosomes with a vacant A site, whose abundance increases during cellular stresses.

## Introduction

Angiogenin was discovered as an angiogenesis factor required for vascularization ^1^ and more recently was found to play roles in other biological processes (reviewed in ^2–4^). Angiogenin is upregulated in cancers, in keeping with the dependence of tumorigenesis on neovascularization ^5–8^. By contrast, missense mutations in angiogenin are correlated with the development of neurodegenerative diseases such as amyotrophic lateral sclerosis (ALS), Parkinson’s disease, and Alzheimer’s disease, highlighting the protein’s key role in neuroprotection ^9–11^.

Angiogenin is a 14-kDa RNase-A family ribonuclease, whose RNase activity is required for angiogenesis, and is characterized by unusual properties ^1,12,13^. First, despite its structural and sequence homology with the highly efficient RNase A-like nucleases, purified angiogenin is at least 10,000-fold less active (k_cat_/K_M_) than RNase A ^12,14–16^. The large difference between the catalytic activities of angiogenin and RNase A correlates with the conformations of the C-terminal tails in protein structures ^15,17,18^. The C-terminus of catalytically competent RNase A adopts a β-sheet conformation that forms a substrate binding surface containing the catalytic histidine corresponding to His114 in angiogenin ^19,20^; (angiogenin numbering is for the 123-aa human protein, excluding the 24-aa signal peptide as in ^21^). By contrast, the C-terminal tail of free angiogenin (residues 116-123), folds into a short α-helix that partially occludes the substrate binding site, consistent with the weak enzymatic activity of purified angiogenin ^14,15^.

Second, angiogenin contrasts the omnipotent RNase A in its high specificity toward cellular tRNAs ^22–25^. Angiogenin cleaves tRNAs at the anticodon stem loop, resulting in 30- to 40-nt-long tRNA fragments and leading to potent inhibition of translation ^22–24,26–30^ and other downstream effects ^31^. The tRNA-cleaving function of angiogenin is also implicated in transgenerational inheritance ^32^, stem cell maintenance ^33^, and astroglial support of motor neurons ^34^. How angiogenin achieves high specificity toward tRNA in cells has been the subject of intense investigation. The catalytic activity of angiogenin substantially increases in cells and cell extracts ^35,36^. Activation of angiogenin during cellular stress coincides with its colocalization with the translational machinery ^37^. These observations suggest that the catalytic function of angiogenin may depend on the translational machinery.

To understand the mechanisms of angiogenesis activation and specificity, we studied how angiogenin interacts with the translation system. We found that the RNase activity of angiogenin is potently activated by the mammalian 80S ribosome. Cryo-EM structures reveal the structural rearrangement of 80S-bound angiogenin into the RNase-A-like conformation, explaining the enzyme’s catalytic activation. Ribosomal positioning of the angiogenin active site next to newly delivered tRNAs underlies the angiogenin’s specificity for the cleavage of tRNA anticodon stem loops. Thus, our work uncovers the 80S ribosome as the activator and specificity factor for angiogenin.

## Results and Discussion

### Angiogenin’s active site is rearranged upon binding 80S ribosome

To investigate how angiogenin inhibits translation, we tested the effect of angiogenin on a mammalian translation system. Consistent with previous reports ^28,29^, translation in different preparations of rabbit reticulocyte lysates (RRL) was inhibited by 70-140 nM human angiogenin (**Fig. 1a**). Addition of angiogenin to RRL resulted in the appearance of 30-40 nt RNA fragments consistent with tRNA cleavage (**Fig. 1b**, **Supplementary Fig. 1a**) as previously reported ^23–25^. Inhibition of angiogenin by RNasin ^38,39^, a commercial version of the protein human ribonuclease/angiogenin inhibitor 1, restored translation and prevented the accumulation of tRNA fragments (**Fig. 1a-b, Supplementary Fig. 1a**), indicating that RNasin may block the interaction of angiogenin with its activating molecule and/or substrate.

**Fig. 1:**
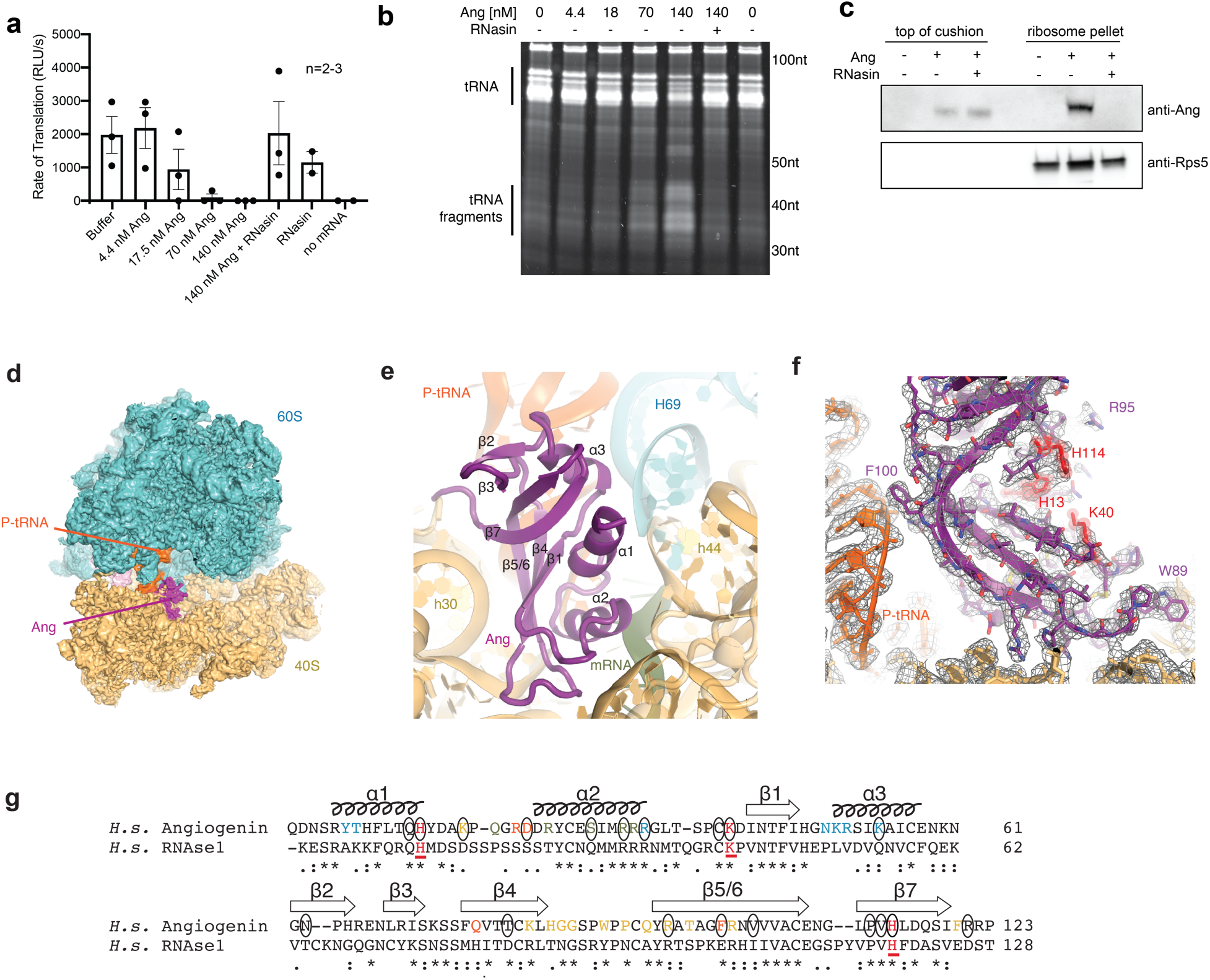
Angiogenin inhibits translation by binding 80S ribosomes. (**a**) Angiogenin inhibits cellular lysate translation (rabbit reticulocyte lysate, RRL) in a concentration-dependent manner, and the inhibition is relieved by RNasin. (**b**) Incubation of RRL with angiogenin for 15 minutes leads to the accumulation of tRNA fragments; fragment accumulation is inhibited by RNasin. (**c**) Angiogenin co-pellets with the ribosome fraction in RRL reactions, except in the presence of RNasin. (**d**) 3.0 Å cryo-EM map of 80S-angiogenin complex. 60S subunit is shown in cyan, 40S subunit in yellow, P-site tRNA in orange, mRNA in green, and angiogenin in magenta. (**e**) Close up view of angiogenin in the A-site of the ribosome with secondary structure elements labeled. (**f**) Cryo-EM density showing well-resolved interactions of angiogenin with P-tRNA and 18S rRNA; the three catalytic residues are shown in red for reference, note the C-terminal residues of angiogenin (118 to 123) and h30 have been omitted for clarity of P-tRNA interactions. Mesh shows 2.8 Å cryo-EM map with B-factor of -50 Å^2^ at 3.5 σ. (**g**) Sequence alignment of human angiogenin with RNase A (RNase1) with secondary structure indicated for angiogenin in the ribosome complex including β-strand 5 joining β-strand 6 (5/6) and β-strand 7. Underlined, red letters indicate catalytic residues conserved between angiogenin and RNase A ^17,18^. Blue letters are angiogenin residues that interact with 28S rRNA, gold letters are angiogenin residues that interact with 18S rRNA, green letters are angiogenin residues that interact with mRNA, and orange letters are angiogenin residues that interact with P-site tRNA in the cryo-EM structure. Circled resides have been found to be mutated in ALS and Parkinson’s patients ^9–11^.

We next tested if angiogenin interacts with the lighter (e.g., individual proteins/RNA) or heavier (e.g., ribosomes) components of the translation system. Sucrose cushion ultracentrifugation revealed that angiogenin is present in the ribosome fraction (**Fig. 1c, Supplementary Fig. 1b**). Addition of RNasin prevents ribosomal association, suggesting that angiogenin function may depend on its interaction with the ribosome.

To explore the interaction of angiogenin with ribosomes, we added angiogenin to 80S•mRNA•tRNA ribosomes and performed structural analysis using cryo-EM. We obtained a 3.0-Å resolution cryo-EM map that shows a well-resolved angiogenin bound to the A site of the small (40S) subunit of the ribosome (**Fig.1d-f, Fig. 2a, Supplementary Fig. 1c**,d). Maximum-likelihood classification of cryo-EM data revealed that among numerous ribosome and tRNA conformations reflecting distinct functional states, angiogenin is bound with the elongation-like ribosomes containing P-site tRNA (**Supplementary Fig. 1c**). Roughly 2200 Å^2^ (∼30%) of angiogenin’s solvent accessible surface area interacts with the ribosome indicating strong and specific binding ^40^. Angiogenin interacts via polar and hydrophobic groups with all elements of the decoding center, including 28S rRNA, 18S rRNA, mRNA, and the adjacent P-tRNA (**Fig. 1e,f**).

**Figure 2.**
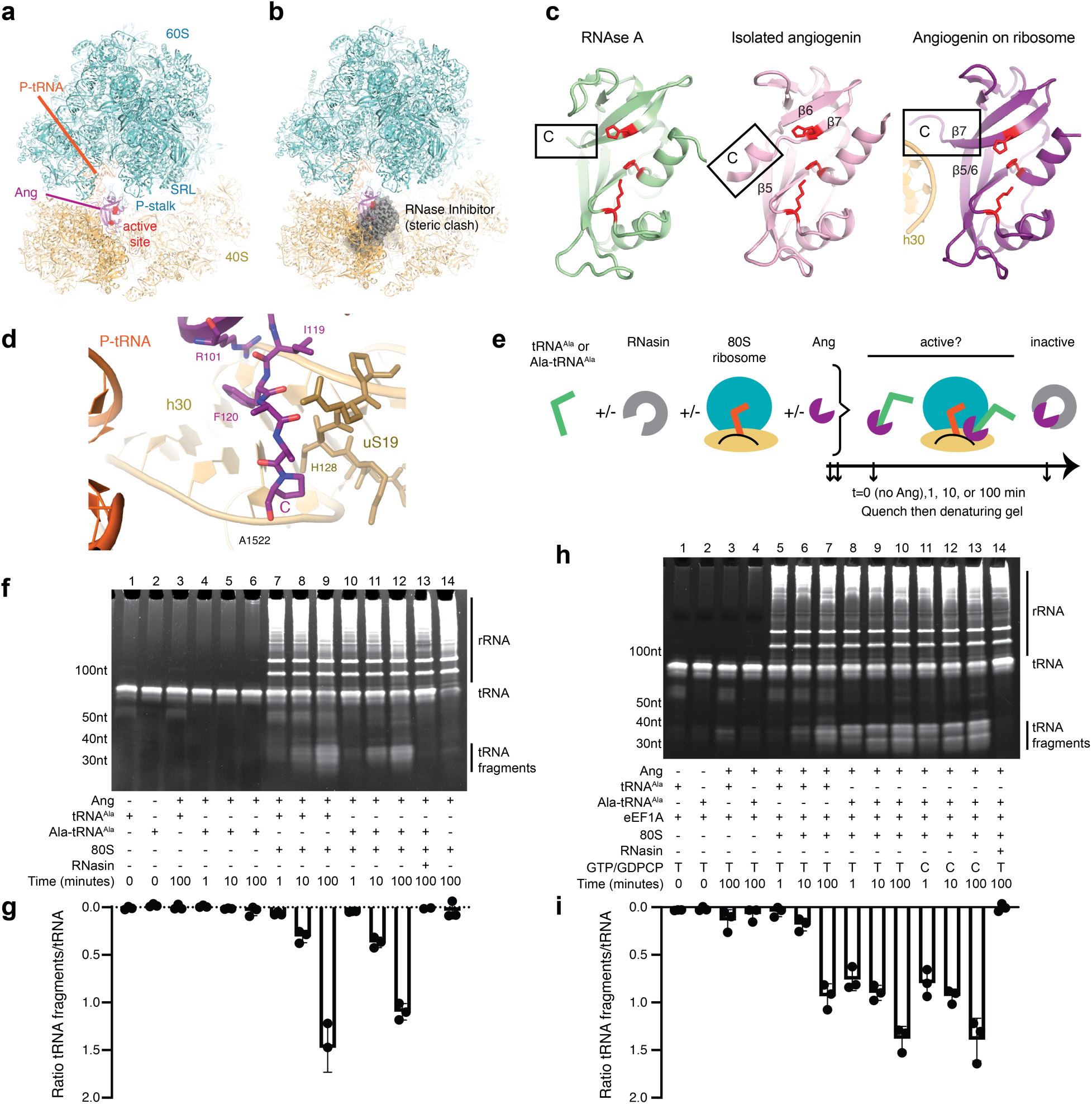
Angiogenin’s RNase is activated upon ribosome binding. (**a**) The catalytic residues (red) of angiogenin (magenta) face the tRNA-delivery path, comprising the tRNA-binding P-stalk and GTPase-activating sarcin-ricin loop (SRL) of the 60S subunit. (**b**) Interaction of angiogenin with the ribosome is incompatible with its interaction with the RNase inhibitor (gray) due to steric clash, explaining angiogenin inhibition by RNasin. Structural alignment compares ribosome-bound angiogenin (this work) with angiogenin bound to human RNase inhibitor (PDB: 1A4Y; ^51^). (**c**) Catalytically active RNase A contains a C-terminal β-strand (left panel; PDB: 5RSA; ^52^), whereas isolated angiogenin has an inhibited α-helical conformation of the C-terminus (middle panel; PDB:5EOP; ^53^). Ribosome-bound angiogenin features a rearranged C-terminus (right panel; this work) similar to that of RNase A. (**d**) Close up view of the C-terminal β-strand of angiogenin stabilized by ribosomal helix 30 (h30) of 18S rRNA, uS19 and β5/6 strand. (**e**) Scheme for assay to measure 80S-stimulated tRNA cleavage by angiogenin is shown with the order of addition of components from left to right. (**f**) Production of tRNA fragments by angiogenin is stimulated in the presence of the 80S ribosome. tRNA cleavage reactions as in (e) were separated by denaturing 15% Urea-PAGE gels stained with SybrGold, one gel is shown (also see **Supplementary Fig. 2**). (**g**) tRNA cleavage by angiogenin is enhanced in the presence of the ribosome. The ratio of signals of tRNA fragments to full length tRNAs (mean +/- standard deviation, n=3). (**h**) Production of tRNA fragments by angiogenin is stimulated in the presence of the 80S ribosome and eEF1A ternary complex with GTP or GDPCP and Ala-tRNA^Ala^ but not with deacyl-tRNA^Ala^ (also see **Supplementary Fig. 2**). (**i**) Ala-tRNA^Ala^ present as a ternary complex is cleaved by angiogenin more efficiently than tRNA alone. The ratio of signals from tRNA fragments to full length tRNA is shown (mean +/- standard deviation, n=3).

Helices α1 and α3 of angiogenin bind the central helix 69 (H69) of 28S rRNA, which docks into the decoding center of the small subunit (**Fig. 1e, Supplementary Fig. 1e**). The positively charged patch of α3, composed of Lys50, Arg51, and Lys54 binds the phosphodiester backbone of H69. These residues are specific to angiogenin (e.g., RNase A features short uncharged side chains), in keeping with their role in ribosome binding (**Fig. 1g**). Interactions with 18S rRNA are predominately formed by strands β4 and β5/6 and their connecting loop (aa 76-101). Trp89 stacks onto 18S nucleotide U627 (U531 in *E. coli*), a key element of the ribosomal decoding center near the mRNA-binding tunnel (**Supplementary Fig. 1f**). Arg33 of α2 bridges the two ribosomal subunits by reaching into the decoding center toward universally conserved nucleotides A3760 at the tip of H69 (corresponding to A1913 in *E. coli* 23S rRNA) and A1825 of the small-subunit helix 44 (h44; A1493 in *E. coli* 16S rRNA) (**Supplementary Fig. 1g**). A patch of polar residues, including Arg24, Ser28, Arg31, and Arg32, reaches toward the A-site mRNA codon and interacts with mRNA nucleotides via electrostatic and stacking interactions (**Supplementary Fig. 1h**-i). Lastly, angiogenin contacts the P-site tRNA by the stacking of Phe100 onto the ribose of the tRNA’s nucleotide 28, while Gln77 reaches the phosphate group of this nucleotide (**Fig. 1f**). Extensive interactions with the ribosomal decoding center, mRNA, and P-site tRNA suggest that angiogenin specifically binds elongating ribosomes whose A site is vacant (i.e., not occupied by tRNA or another ligand) due to ribosome stalling or cellular stress ^41–44^. The ribosome-binding activity of angiogenin may account for loss-of-function mutations that were not previously explained by angiogenin’s cell-receptor binding ^45^, nuclear localization ^46^, or other processes ^47–50^. For example, many ribosome interactions involve angiogenin’s residues whose mutations are associated with ALS and Parkinson’s disease (**Fig. 1g**, ^9–11^).

Angiogenin’s conformation is altered by ribosome binding, suggesting a mechanism for the catalytic activation of the enzyme (**Fig. 2**). The activity of free angiogenin is several orders of magnitude lower than that of RNase A (**Fig. 2c, left panel**) ^12,14–16^, consistent with the inhibitory α-helical conformation of the C-terminal tail of angiogenin ^17^, which occludes substrate binding (**Fig. 2c, middle panel**). In the 80S•angiogenin ribosome complex, the C-terminus of angiogenin is shifted by more than 15 Å and rearranged to extend β-strand 7 (**Fig. 2c, right panel**), similar to that of free RNase A (**Fig. 2c, left panel**). In this conformation, the C-terminus is stabilized by interactions with the minor groove of 18S rRNA helix 30 (h30) and the neighboring β-strands 5 and 6, which are reorganized into a single longer stand that we term β5/6 (**Fig. 2c**). Here, Phe120 of angiogenin packs between A1522 of h30 and Arg101 of β5/6 docked at the phosphate group of C1237, while the neighboring Ile119 binds near the C-terminus of uS19 (**Fig. 2d**). In this rearranged angiogenin conformation, catalytic residues His13 and His114 are positioned to catalyze substrate cleavage as in RNase A (**Fig. 2c**). The active site is exposed in the intersubunit space and is fully accessible to tRNA substrates (**Fig. 2a**).

### 80S ribosome stimulates angiogenin’s tRNA cleavage activity

To test if the ribosome directly activates angiogenin’s RNase activity, we compared the ability of angiogenin alone or in complex with the elongation-like 80S ribosome to cleave purified tRNA^Ala^—an angiogenin substrate known to generate inhibitory tRNA fragments ^22^ (**Fig. 2e-g, Supplementary Fig. 2**). Almost no tRNA^Ala^ cleavage was detected in the presence of 10 nM angiogenin at 100 minutes (**Fig. 2f-g, lane 3**), consistent with previous reports of poor activity of isolated angiogenin ^12,14–16^. By contrast, within one minute of adding tRNA^Ala^ to a reaction containing 10 nM angiogenin with 20 nM 80S ribosome complex, ∼35-nt tRNA fragments began to accumulate; and after 100 minutes, we detected >100-fold accumulation of cleavage products (**Fig. 2f-g, lane 7 and 9**). RNasin prevented tRNA cleavage in the presence of the 80S complex (**Fig. 2f-g, lane 13**), consistent with RNasin’s ability to block angiogenin’s binding to the ribosome (**Fig. 1c**). Indeed, angiogenin’s α1 and α3, which interact with the ribosomal H69 and P-site tRNA, are occluded in the complex of angiogenin with human RNase inhibitor, an natural version of RNasin ^51^; and the angiogenin•RNase inhibitor complex is incompatible with ribosome binding due to a severe clash of RNasin with the 40S subunit (**Fig. 2b**). Thus, the 80S ribosome potently stimulates tRNA cleavage into ∼35-nt fragments by angiogenin, consistent with the cryo-EM structure described above.

We next asked whether the translation elongation factor eEF1A, which delivers aminoacyl-tRNA to the ribosome during translation elongation in cells, may present tRNA to the 80S•angiogenin complex for cleavage. The tRNA cleavage assay revealed that eEF1A further stimulated tRNA cleavage of amino-acylated tRNA^Ala^ by the ribosome-bound angiogenin, resulting in the 10- to 100-fold higher accumulation of tRNA fragments within 1 minute than that without eEF1A (**Fig. 2h-i, lane 8 vs. Fig. 2f-g, lanes 7 and 10**). By contrast, eEF1A did not stimulate deacyl-tRNA cleavage, consistent with the inability of eEF1A to bind deacyl-tRNA ^54^ (**Fig. 2h-i, lane 5**). Although GTP hydrolysis on eEF1A is required for elongation ^54^, eEF1A stimulated amino-acylated tRNA cleavage similarly in the presence of either GTP or non-hydrolyzable GTP analog, GDPCP (**Fig. 2h-i, lane 8 vs. lane 11**). These results suggest that eEF1A binding to the ribosome results in optimal presentation of tRNA to the angiogenin catalytic site, but hydrolysis of GTP by eEF1A is not required for tRNA cleavage.

### Cryo-EM structure of the 80S•angiogenin complex with eEF1A and tRNA

To understand how tRNA is delivered and positioned for cleavage by ribosome-bound angiogenin, we used cryo-EM to analyze the 80S•angiogenin complex supplemented with eEF1A•Ala-tRNA^Ala^•GDPCP ternary complex. Maximum-likelihood classification of the cryo-EM dataset revealed three types of ribosome complexes bound with angiogenin (**Supplementary Fig. 3a**): (1) 80S•angiogenin as described above, at an improved resolution of 2.8 Å (**Fig. 1f**; also see Methods describing a minor fraction of partially rotated ribosomes with P-site tRNA); (2) 80S•angiogenin with eEF1A and Ala-tRNA^Ala^ (**Fig. 3a, Supplementary Fig. 3c**,f**,h,i**); and (3) 80S•angiogenin with tRNA^Ala^ (**Fig. 3b, Supplementary Fig. 3d**,g). The tRNA substrate-bound ribosome particles represent only a fraction (<20%) of angiogenin-bound ribosomes, consistent with transient interactions between the incoming tRNA and activated enzyme in the multiple-turnover reaction. In both complexes containing tRNA^Ala^, the anticodon loop was unresolved, suggesting that the maps contain the products of fast tRNA cleavage (**Supplementary Fig. 3c**-d). Indeed, PAGE analysis of the cryo-EM reaction shows that tRNA^Ala^ is cleaved (**Supplementary Fig. 3e**).

**Figure 3.**
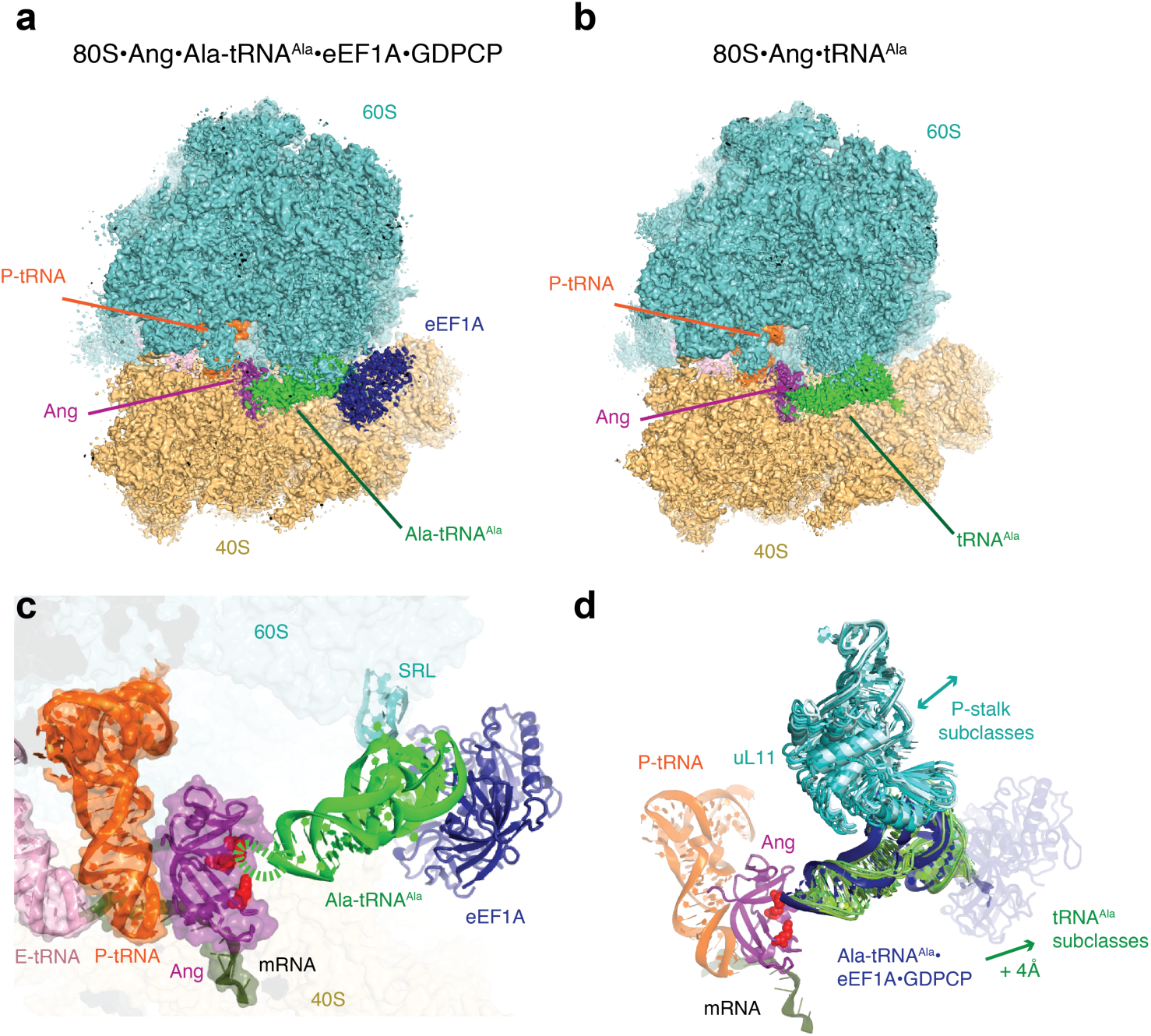
Cryo-EM structures of 80S•angiogenin complexes formed with eEF1A•GDPCP and tRNA. (**a**) Cryo-EM map of 80S•angiogenin bound with eEF1A•Ala-tRNA^Ala^, colored as in **Fig. 1** with the addition of green for Ala-tRNA^Ala^ and dark blue for eEF1A. (**b**) Cryo-EM map of 80S•angiogenin bound with tRNA^Ala^. (**c**) The anticodon stem of Ala-tRNA^Ala^ (green) in the ternary complex is placed in the vicinity of the angiogenin active site (red), and the anticodon loop is not resolved due to cleavage and/or mobility (dashed green line). (**d**) Different positions for tRNA^Ala^ among five different cryo-EM maps suggest tRNA mobility near angiogenin. The 60S P-stalk (shades of cyan) and tRNA (shades of green) exhibit different positions between classes indicative of ∼8 Å movement.

In the structure with eEF1A and Ala-tRNA^Ala^, the phosphate of the last modeled tRNA nucleotide of the anticodon stem (U38) is placed near the active site of angiogenin, consistent with catalytic cleavage near this position (**Fig. 3c**). Indeed, based on the preference of angiogenin to cleave between a pyrimidine and an adenosine residue ^55^, the anticodon loop of tRNA^Ala^ (^32^TTAGCAT^38^) can be cleaved at positions 33 and 36, although our current work does not distinguish whether one or both or alternate sites have been affected. Cryo-EM density for tRNA^Ala^ is less well resolved than angiogenin (**Supplementary Fig. 3c**) indicating heterogeneity in the conformation or composition (e.g., various cleavage positions) of the tRNA. Nevertheless, the tRNA canonical L-shaped structure is clearly visible (**Supplementary Fig. 3f**) except for the anticodon loop (**Supplementary Fig. 3c**), indicating that cleavage does not disrupt the overall fold of tRNA. Two major contacts hold the nicked tRNA in place: the elbow interacts with the 28S rRNA of the dynamic P stalk (at nts G19 of tRNA^Ala^ and A2009 of 28S rRNA), similarly to the interaction observed during mRNA decoding ^56^; and the acceptor arm of tRNA binds eEF1A, which is docked on the 40S ribosomal subunit ∼10 Å away from the sarcin-ricin loop (SRL) of the large subunit (**Fig. 3c, Supplementary Fig. 3f**,i). This placement of eEF1A away from the SRL—the site of GTPase activation in mRNA-decoding complexes ^56,57^—agrees with our observation that GTP hydrolysis is not required for tRNA cleavage by ribosome-bound angiogenin.

In comparison with the eEF1A-bound tRNA (dark blue), deacyl tRNA (green) shifts outward by more than 4 Å. In the 80S•angiogenin•tRNA^Ala^ structure, the cleaved tRNA is placed slightly differently than in the eEF1A-containing map (**Fig. 3b,d; Supplementary Fig. 3d**,g). While the anticodon stem of tRNA is placed near angiogenin, the elbow is held further out by the P stalk. Subclassification of the cryo-EM density corresponding to this complex suggests several positions of P stalk and the tRNA, differing by up to 8 Å at the tip of the acceptor arm (**Fig. 3d, Movie S1 and S2**). The flexible tethering of tRNA by the P stalk may allow access of different nucleotides within the anticodon loop to the angiogenin’s active site, accounting for cleavage of various or possibly all tRNA species ^26,30^. The 3′-CCA end is close to or docked at 18S rRNA helix 5 (h5) near residue A50 (**Supplementary Fig. 3g, Movie S2**). This interaction is sterically incompatible with binding of eEF1A, suggesting that this structure may represent a deacylated tRNA that initially bound the ribosome without eEF1A. Thus, the cryo-EM structures with nicked tRNA^Ala^ show how ribosome-bound angiogenin can cleave tRNAs whether or not they are delivered by eEF1A, consistent with our biochemical observations (**Fig. 2f-i**).

## Conclusions

Elucidating the function and therapeutic potential of angiogenin requires an understanding of the mechanism of angiogenin activation. Our work shows that angiogenin is activated by ribosomes with a vacant A site, which were shown to accumulate during cellular stress ^41–44^. Ribosome binding rearranges the active site of angiogenin, directing the catalytic residues toward incoming tRNA substrates. In cells, tRNAs are commonly bound with eEF1A (and are aminoacylated) or are deacylated (e.g. during cellular stress). We show that both aminoacylated and deacylated tRNAs are substrates for ribosome-bound angiogenin. Incoming tRNAs are held in place by the ribosomal P-stalk, with the anticodon loop presented at the angiogenin active site resulting in the rapid cleavage of tRNA. This structural mechanism accounts not only for the activation of angiogenin, but also for the substrate specificity provided by the ribosome, whose P stalk is tuned for tRNA delivery. Angiogenin was proposed to also cleave the 3′ CCA ends of tRNA ^58^, however recent genetic and tRNA sequencing analyses argue that angiogenin does not have the CCA-cleaving activity and that another RNase must be responsible for the 3′ cleavage ^30^. In our structures, the CCA ends of tRNA bind >70 Å away from angiogenin’s active site (**Fig. 3**), in keeping with angiogenin’s selective cleavage of the anticodon loops. Notably, the resulting nicked tRNA is held together by stacking interactions at tRNA stems and retains the overall tertiary structure of tRNA (**Fig. 3**), as opposed to separated single-stranded tRNA “fragments” that are often implicated in downstream functions. Thus, angiogenin-mediated tRNA cleavage could inhibit translation by depleting functional tRNAs and/or by producing non-productive stalled complexes of tRNA-like cleavage products with tRNA-interacting molecules, such as ribosomes and tRNA-binding enzymes. Future work will investigate the downstream effects of the tRNA-like cleavage products and will explore therapeutic avenues to targeting the ribosome-driven activation of angiogenin.

## Methods

### Preparation of angiogenin, tRNA and eEF1A

Recombinant human angiogenin (catalog number 265-AN/CF) was purchased from R&D Systems, a Bio-Techne Company (Minneapolis, MN). The lyophilized stock was resuspended as recommended by the manufacturer in phosphate buffered saline at 1 mg/ml or 0.1 mg/ml. The molar concentrations were derived from A280 measurements using the calculated extinction coefficient from https://web.expasy.org/protparam/.

Aliquots were flash frozen in liquid nitrogen and stored at -80°C. Two independently ordered lots were used in biochemical and cryo-EM experiments, and both gave similar results (Lot DK0420031 and Lot DK0421061).

### tRNA^Ala^

Yeast tRNA^Ala^ (anticodon AGC) was produced by *in vitro* transcription as follows: the gene encoding tRNA^Ala^ (anticodon AGC) and preceded with the T7 RNA polymerase promoter was synthesized and cloned into pUC57 between the EcoRI and BamHI sites by GeneScript USA Inc. (Piscataway NJ). The T7 promoter and tRNA^Ala^ encoding region was then PCR amplified with primers (Integrated DNA Technologies, Inc, Coralville IA) encoding the T7 promoter (5’-GAATTCTAATACGACTCACTATAGGGCGTG) and the 3’ end of the tRNA including the non-templated CCA of mature tRNA (5’-TGGTGGACGAGTCCGG-3’). The PCR product was used as a template for run-off *in vitro* transcription using T7 polymerase as previously described ^59^. The *in vitro* transcribed tRNA (IVT tRNA) was folded by heating to 65°C and slowly cooling to room temperature for 30 minutes in Refolding buffer (20 mM KOAc pH 7.5, 20 mM MgCl_2_). tRNA purity was confirmed by PAGE and concentration was determined using the absorbance at 260 nm (1800 pmol/ml per A260 (1cm)). The stock solution was stored at -80°C.

tRNA^Ala^ was charged with alanine using yeast S100 extract from the protease-deficient strain BJ2168. To prepare S100 extract, 6 L of BJ2168 cells were grown until OD600 of1.5 in YPD, pelleted and then flash frozen. The frozen cells were ground in a ball-mill grinder (Retsch GmbH, Hann Germany) and resuspended in 20 ml of S100 Lysis Buffer (10 mM KHPO_4_ pH 7.5, 1x Mini protease inhibitor cocktail tablet (Roche, Basel, Switzerland)) for 1 hour. This and all subsequent steps took place at 4°C. The lysate was clarified by centrifugation at 20,000xg for 0.5 hours (Beckman Coulter, Brea, CA). The ribosomes were pelleted from the clarified lysate by ultracentrifugation in a Ti-70 rotor at 100,000xg for 3 hours. The top 80% of the post-ribosomal supernatant was collected and incubated with S100-Lysis-buffer-pre-washed DEAE cellulose resin (Sigma-Aldrich, St. Louis, MO, 2.5 g dry weight before washing) for 0.5 hours. The resin was recovered and washed twice with 10 column volumes of S100 Lysis Buffer on a gravity flow column (Bio-Rad Laboratories Inc., Hercules, CA). The S100 extract was eluted from DEAE resin with 5 ml of S100 Elution Buffer (250 mM KHPO_4_ (pH 6.5),1x Mini protease inhibitor cocktail tablet (Roche, Basel, Switzerland)) and aliquoted and flash-frozen in liquid nitrogen. Folded IVT tRNA^Ala^ (77 μM) was incubated with 1 mM L-alanine and 20% *S. cerevisiae* S100 extract in 1xCharging Buffer (50 mM Hepes-KOH pH 7.5, 10 mM KCl, 50 mM MgCl_2_, 30 μM ZnCl_2_, 12 mM ATP, 1 mM DTT) at 25°C for 20 hours. The charged tRNA (Ala-tRNA^Ala^) was phenol extracted twice, chloroform:isoamyl (24:1) extracted twice, and then ethanol precipitated in the presence of NaOAc pH 5.2. The resulting pellet was washed with cold, 70% ethanol. The precipitated Ala-tRNA^Ala^ was resuspended in 2 mM NaOAc pH 5.2. Aminoacylated-tRNA was further separated from excess ATP by passing through a Sephadex G-25 column in 2 mM NaOAc pH 5.2. The concentration of sample was determined using the absorbance at 260 nm (1800 pmol/ml per A260 (1cm) unit) and the extent of aminoacylation was determined by acidic PAGE ^60^ and was >90%. The Ala-tRNA^Ala^ solution was stored at -80°C prior to use.

### Purification of eEF1A

Yeast eEF1A was purified from *Saccharomyces cerevisiae* BJ2168 with some modification ^61^. Yeast eEF1 is >80% sequence-identical with rabbit eEF1A and can fully substitute the function of mammalian eEF1A in a mammalian translation system ^62^. To obtain endogenous eEF1A, 6.5 ml of flash frozen BJ2168 cells were ground in a liquid nitrogen cooled ball mill (Retsch GmbH, Mann, Germany) and were resuspended in 10 ml of Buffer A (20 mM Tris-HCl pH 7.5, 0.1 mM EDTA, 25% Glycerol, 100 mM KCl, 1 mM DTT, 1x Mini protease inhibitor cocktail tablet (Roche, Basel, Switzerland), 2 mM PMSF (Sigma-Aldrich, St. Louis, MO) and stirred for 1 hour at 4°C. The lysate was clarified by centrifugation at 31,745xg in a JA20 rotor (Beckman Coulter, Brea, CA) for 15 minutes at 4°C and the soluble portion was retained. This clarification step was repeated twice. The clarified lysate was loaded onto a pre-washed/pre-equilibrated CM Sepharose HiTrap column (5 ml, GE Healthcare, Chicago, IL) on an Äkta explorer FPLC (GE Healthcare, Chicago, IL). The column was washed with 12 column volumes (CV) of Buffer A and then eluted with a 0-100% gradient of Buffer B (20 mM Tris pH 7.5, 0.1 mM EDTA, 25% Glycerol, 1 M KCl, 1 mM DTT, 1x Mini protease inhibitor cocktail tablet (Roche, Basel, Switzerland), and 2 mM PMSF) over 10 CV. Elution profile and SDS-PAGE analysis revealed that eEF1A eluted at 23.4% gradient of Buffer B. The peak fractions were diluted to the final KCl concentration of 50 mM with Buffer C (20 mM HEPES pH 7.5, 0.1 mM EDTA, 25% Glycerol, 1 mM DTT, 1x Mini protease inhibitor cocktail tablet (Roche, Basel, Switzerland) and filtered (0.2 µm pore size). Then the solution was injected onto a prewashed SP Sepharose HiTrap column (5 ml, GE Healthcare, Chicago, IL). The column was washed with 20 CV of Buffer D (20 mM HEPES-KOH pH 7.5, 0.1 mM EDTA, 25% Glycerol, 50 mM KCl, 1 mM DTT, 1x Mini protease inhibitor cocktail tablet (Roche, Basel, Switzerland)) and then was eluted by a stepwise gradient with Buffer E (20 mM HEPES pH 7.5, 0.1 mM EDTA, 25% Glycerol, 1 M KCl, 1 mM DTT, 1x Mini protease inhibitor cocktail tablet (Roche, Basel, Switzerland)) as follows: 15% Buffer E for 5 CV, 30% Buffer E for 5 CV, 50% Buffer E for 5 CV, and 100% Buffer E for 5 CV. The eEF1A fraction eluted at ∼30% Buffer E and was exchanged to Buffer F (20 mM HEPES pH 7.5, 0.1 mM EDTA, 25% Glycerol, 100 mM KCl, 2 mM MgCl2, 1 mM DTT, 1x Mini protease inhibitor cocktail tablet (Roche, Basel, Switzerland)) using an Amicon Ultra centrifugal filter unit with MWCO of 10kDa (MilliporeSigma, Burlington MA). The protein was further purified by gel filtration chromatography on a Superdex 75 16/60 column with Buffer F. The eEF1A-containing peak was identified by 12%SDS-PAGE, pooled, concentrated, and stored at -20°C.

### Translation in rabbit reticulocyte lysates

Rabbit reticulocyte lysates (RRL, Promegga^TM^ Corp., Madison, WI) were used to study the inhibition of translation by angiogenin. We monitored translation of an mRNA encoding nanoluciferase ^63^ in nuclease-treated RRL (Promega^TM^ Corp., Madison, WI). In these experiments, 50% (final concentration) nuclease-treated RRL (Promega^TM^ Corp., Madison, WI) was supplemented with 0.02 mM amino acids (Promega^TM^ Corp., Madison, WI), 1% Nano-Glo® nanoluciferase substrate (Promega^TM^ Corp., Madison, WI) and the indicated concentration of angiogenin (R&DSystems, Minneapolis, MN) or 1xPBS with or without 0.8 U/μl rRNasin (Promega^TM^ Corp., Madison, WI) for 15 minutes at 30^°^C, similarly to the 15-minute angiogenin preinucbation described previously ^24^. Next, 10 ng/μl (final concentration) 6xHis-Nanoluciferase mRNA flanked with the 5’- and 3’-UTR from rabbit beta-globin ^64^ was added, forming a 10 μl reaction mixture, and translation was followed over time by measuring the luminescence of Nanoluciferase using an Infinite® m1000 PRO or a Spark® microplate reader (Tecan Trading AG, Switzerland) for 15 minutes at 30^°^C. The slope of the luminescence signal between 200 to 300 s (initial burst of nanoluciferase synthesis for an uninhibited reaction) was measured in Prism 8 (GraphPad Software, LLC) for each condition across three separate experiments and is plotted as “Rate of Translation (RLU/s)” in **Fig. 1a**. For **Fig. 1b**, the 9 μl reaction mixtures containing RRL and angiogenin prior to mRNA addition were stopped at 15 minutes by 5 volumes of 8M Urea Loading Buffer (8M Urea, 2 mM Tris-HCl pH 8, 20 mM EDTA). When all reactions were completed, the reactions were heated at 95°C for 5 minutes, spun down, vortexed and 5 μl was loaded onto a 15% pre-cast Novex Urea-TBE gel (Thermo Fisher Scientific Corp, Waltham MA) and run at 180V for 60 minutes alongside Small RNA Marker (Ab Nova GmbH, Taipei City, Taiwan). The gels were stained in 1xTBE with 1:20,000 SYBR^TM^ Gold nucleic acid stain (Thermo Fisher Scientific Corp, Waltham MA) for 30 minutes and imaged on a ChemiDoc MP system (Bio-Rad Laboratories Inc., Hercules, CA) using the UV light mode for SYBR^TM^ Gold.

### Association of angiogenin with ribosome fraction

Separation of RRL translation reactions into supernatant and ribosome pellet fractions was performed in RRL left untreated with nuclease, so that it contained endogenous mRNA. Translation reactions (50 μl, containing 50% RRL (Green Hectares; https://greenhectares.com/), 30 μM hemin (Sigma-Aldrich, St. Louis, MO), 75 mM KCl, 0.5 mM MgCl_2_, 1 mM ATP, 0.2 mM GTP, 2.1 mg/ml creatine phosphate, 30 μM of L-amino acid mix, 15 mg/ml yeast tRNA, 6 mM βME, all final concentrations) were assembled adding as the last component (a) 1.4% 1xPBS or (b) 1 μM angiogenin (in 1xPBS) (R&DSystems, Minneapolis, MN) or (c) 1 μM angiogenin preincubated for 5 minutes with 8 U/μl (final concentrations) rRNasin (Promega^TM^ Corp., Madison, WI). Reactions were performed in a heat block at 37°C. After 5 minutes, reactions were terminated by addition of 100 μl of ice-cold 1×10 Buffer (50 mM Tris-Acetate pH 7.0, 100 mM KOAc, 10 mM Mg(OAc)_2_) with 10 mM βME. The reactions were applied to 15% w/v sucrose cushion in 1×10 Buffer with 10 mM βME) and spun in an TLA-110 rotor in an Optima MAX-TL ultracentrifuge (Beckman Coulter, Brea, CA) at 408,800xg (avg) for 1 hour at 4°C. The top third (1 ml) of the sucrose cushion was collected as the supernatant fraction and 2 μl were loaded per lane on the Western blots shown in **Fig. 1c** and **Supplementary Fig. 1b**. The ribosome pellet was washed in 1×10 Buffer with 10 mM βME and then was resuspended in 40 μl of 1×10 Buffer with 10 mM βME using a microstirbar and pipetting. The A_260_ of the ribosome fractions was measured and 10 OD units were loaded per lane on the Western blots shown in **Fig. 1c** and **Supplementary Fig. 1b**.

### Western Blotting

Samples alongside Precision Plus Protein Dual Color Standard ladder (Bio-Rad Laboratories Inc., Hercules, CA) were run on 4-20% Mini-Protean® TGX^TM^ Precast Protein gels (Bio-Rad Laboratories Inc., Hercules, CA) at 120 V for 1 hour in 1xTris-Tricine-SDS running buffer. The gels were transferred to PVDF membrane, 0.2 micron pore size (Bio-Rad Laboratories Inc., Hercules, CA) in 1xTris-Glycine buffer with 20% methanol. The membranes were blocked in 1xPBS + 0.1%Tween (1xPBST) buffer with 6% Blotto non-fat milk powder (Santa Cruz Biotechnologies, Dallas, TX). Following 4 washes in 1xPBST shaking for 5 minutes each, the membranes were incubated with primary antibody either 1:500 mouse anti-angiogenin monoclonal antibody C-1 (sc-74528, Santa Cruz Biotechnologies, Dallas, TX) or 1:500 mouse anti-ribosomal protein s5 monoclonal antibody (A-8) (sc-390935, Santa Cruz Biotechnologies, Dallas, TX) in 1XPBST with 6% milk powder for 1 hour at room temperature or overnight at 4°C. Following 4 washes shaking in 1xPBST for 5 minutes each, the membranes were incubated with secondary antibody 1:10,000 goat anti-mouse IgG (H+L)-HRP conjugate (62-6520, Thermo Fisher Scientific Corp, Waltham MA) in 1xPBST + 6% milk powder for 30 minutes. Following 4 washes in 1xPBST for 5 minutes each, the blot was developed with SuperSignal^TM^ West Femto Maximum Sensitivity Substrate (Thermo Fisher Scientific Corp, Waltham MA) and imaged on a ChemiDoc MP system (Bio-Rad Laboratories Inc., Hercules, CA) using the chemiluminence setting for Western imaging and colorimetric setting to capture the ladder.

### Cryo-EM samples of mammalian 80S•angiogenin, with and without eEF1A•GDPCP•Ala-tRNA^Ala^ ternary complex

*Oryctolagus cuniculus* (European rabbit) 80S ribosomes with angiogenin were prepared with or without a ternary complex of eEF1A, Ala-tRNA^Ala^, and GDPCP. The ribosomal subunits were purified from nuclease-untreated rabbit reticulocyte lysate (RRL) as described ^65^. Rabbit 60S subunits (2 µM) were pre-incubated for 5 minutes at 30°C in 1×10 Buffer (20 mM Tris-acetate (pH 7.0), 100 mM KOAc, 10 mM Mg(OAc)_2_). Meanwhile, 40S subunits were preincubated with a leaderless mRNA encoding for the start codon in the P-site and the UUC Phe codon in the A-site (CCAC-AUG-**<u>UUC</u>**-CCC-CCC-CCC-CCC-CCC-CCC, Integrated DNA Technologies, Inc, Coralville IA) for 5 minutes at 30°C in the same buffer. *E. coli* tRNA^fMet^ (Chemical Block Ltd., Moscow, Russia) was added to 40S such that final concentrations were 1.2 µM 40S, 40 µM mRNA, and 2.4 µM tRNA^fMet^. The 40S and 60S reactions were mixed 1:1 and incubated for 5 minutes at 30°C to form the 80S ribosome with tRNA^fMet^ and AUG codon in the P site. Next, angiogenin was added to 4 µM while diluting the 80S complex by half in 1×10 Buffer and incubated for 5 minutes at 30°C. The resulting 80S-angiogenin complex (containing 0.5 µM 60S, 0.3 µM 40S, 10 µM mRNA, 0.6 µM tRNA^fMet^ and 4 µM angiogenin) was flash frozen in liquid nitrogen, stored at -80°C and was thawed immediately prior to grid plunging. A ternary complex of eEF1A, Ala-tRNA^Ala^, and GDPCP was assembled in 1×10 buffer by preincubation of eEF1A and GDPCP (Jena Bioscience, Jena, Germany) for 5 minutes at 30°C followed by addition of Ala-tRNA^Ala^ for 1 minute at 30°C. The resulting ternary complex reaction (containing 24 µM eEF1A, 4 µM Ala-tRNA^Ala^, and 1 mM GDPCP) was also flash frozen in liquid nitrogen and was stored at -80°C.

For both 80S•angiogenin complex and 80S•angiogenin•ternary-complex, Quantifoil R2/1 holey-carbon grids coated with a thin layer of carbon (Electron Microscopy Service, Hatfield, PA) were glow discharged with 20 mA current with negative polarity for 30 s (with ternary complex) or 60 s (without ternary complex) in a PELCO easiGlow glow discharge unit. The Vitrobot Mark IV (Thermo Fisher Scientific Corp, Waltham MA) was pre-equilibrated to 5°C and 95% relative humidity. For 80S-angiogenin complex, 2.5 µl of the 80S•angiogenin reaction was applied to grid, incubated for 10 s, blotted for 4 s, and then plunged into liquid-nitrogen-cooled liquid ethane. For the 80S•angiogenin•ternary complex reaction, 1.5 µl of 80S•angiogenin was mixed with 1.5 µl of ternary complex in an ice-cold tube with ice-chilled tips and was applied as quickly as possible to the grid or with a 30 s delay. The solution was incubated on the grid for 10 s, blotted for 4 s, and then plunged into liquid-nitrogen-cooled liquid ethane. The total elapsed time from mixing 80S•angiogenin with ternary complex to cryo-plunging was either 25 s or 60 s. 15% TBE-urea gel electrophoresis analysis of a time course of the same reactions (80S•angiogenin and ternary complex) showed that cleavage of Ala-tRNA^Ala^ nearly saturates within 25 s (**Supplementary Fig. 3e**) thus the 25 s time point was chosen for cryo-EM analysis.

### Electron Microscopy

Data were collected on a Talos electron microscope (Thermo Fisher Scientific Corp, Waltham MA) operating at 200 kV and equipped with a K3 direct electron detector (Gatan Inc.) targeting 0.5 to 1.8-μm underfocus. Data collection was automated using SerialEM ^66^ using beam tilt to collect multiple movies (e.g. 5 movies per hole at 4 holes) at each stage position ^67^. The rabbit 80S•angiogenin dataset had 1015 movies, at 20 frames per movie at 1.1 e^-^/Å^2^ per frame for a total dose of 29.6 e^-^/Å^2^ on the sample and yielding 107,236 particles. The rabbit 80S•angiogenin•ternary complex dataset had a total of 20 frames per movie, with 1.55 e^-^/Å^2^ per frame for a total dose of 30.9 e^-^/Å^2^ on the sample, comprising 5,044 movies yielding 273,774 particles. The movies were aligned during data collection using IMOD ^68^ to decompress frames, apply the gain reference, and to correct for image drift and particle damage and bin the super-resolution pixel by 2 to 0.87 Å in the final image sums.

### Cryo-EM data classification and map refinement

CTF determination, reference-free particle picking, and stack creation were carried out in cisTEM ^69^. Particle alignment and refinement was carried out in Frealign 9.11 and cisTEM ^69,70^. Box size for both datasets was 608x608x608. To speed up processing, binned image stacks (e.g. 8×, 4× or 2×) were prepared using resample.exe part of the Frealign v9.11 distribution. The initial model for particle alignment of 80S maps was EMD-4729, https://www.ebi.ac.uk/emdb/EMD-4729 ^71^, which was downsampled to match the 8xbinned stack and low-pass filtered to 30 Å, using EMAN2 ^72^. Two rounds of mode 3 search alignment to 30 then 20 Å were performed using the 8xbinned stack. Next, rounds of mode 1 refinement were run with the 4x binned and eventually the 2x binned stack as we gradually added resolution shells to 8 Å.

For the 80S•angiogenin dataset, focused 3D maximum-likelihood classification into 6 classes (using the high-resolution limit of 12 Å and the 4xbinned stack, 30 rounds) with a 45-Å spherical mask covering the A site of the ribosome (x, y, z = 225.405, 278.628, 242.574) resolved different ribosome states including two angiogenin-bound classes. The particles from these two classes were extracted using merge_classes.exe setting a threshold of class occupancy of 0.50 and score of 0. The resulting stack of 45,850 particles was independently refined starting from the EMD-4729 starting model until the high-resolution limit of 5 Å and was further improved by one round of per particle CTF refinement to yield the final map at 3.04 Å resolution.

For the 80S•angiogenin•ternary complex dataset, focused 3D maximum-likelihood classification into 8 classes (using the high-resolution limit of 15 Å and the 8xbinned stack, 50 rounds) with a 50-Å spherical mask covering the A site and the GTPase-activating center of the ribosome (x, y, z = 225.756, 296.851, 200.364) resolved different ribosome states (**Supplementary Fig. 3a**) including the three 80S•angiogenin classes, one 80S•angiogenin•tRNA^Ala^ state. The three 80S•angiogenin classes were merged yielding a 101,572-particle stack, independently refined and improved with both per-particle CTF refinement followed by beam-tilt refinement to yield a map at 2.81 Å resolution (high resolution limit for refinement was 4 Å). The 80S•angiogenin•tRNA^Ala^ particles were extracted and further separated using focused 3D maximum-likelihood classification into 6 classes (using the high-resolution limit of 15 Å and the 8xbinned stack, 100 rounds) using a 60-Å mask around A-site and GTPase-activating center and encompassing eEF1A (x, y, z = 253.211, 303.049, 190.051). This classification yielded five 80S•angiogenin•tRNA^Ala^ classes and one 80S•angiogenin•Ala-tRNA^Ala^•eEF1A•GDPCP state (**Supplementary Fig. 3a**). The five 80S•angiogenin•tRNA^Ala^ classes were merged yielding a 17,593 particle stack, independently refined and improved with per-particle CTF refinement followed by beam-tilt refinement to yield a map at 3.09 Å resolution (high resolution limit for refinement was 5 Å). The 80S•angiogenin•Ala-tRNA^Ala^•eEF1A•GDPCP particles were independently extracted yielding a 3,604 particle stack that once refined and beam-tilt corrected yielded a map at 3.67 Å resolution (high resolution limit for refinement was 9 Å). Finally, the particles from the rotated 40S class were extracted yielding a 36,105 particle class that was further separated using focused 3D maximum-likelihood classification into 6 classes (using the high-resolution limit of 15 Å and the

8xbinned stack, 50 rounds) and a 50-Å mask around the head of 40S subunit and encompassing the three tRNA binding sites (x, y, z = 194.690, 263.751, 232.504). This classification yielded three rotated, hybrid-state classes with either one or two tRNAs (P/E or A/P and P/E) and without angiogenin; two classes with partial subunit rotation (∼8.5*°* 40S body rotation and ∼0*°* head rotation) and with classical-state-like tRNAs in the E/E and P/P sites and angiogenin in the A site; and one non-rotated class with two classical-state tRNAs and weak angiogenin density. Angiogenin density was lower-resolution in the partially subunit-rotated classes than in the main maps described above and angiogenin interacts predominantly with H69.

### Model building and refinement

Cryo-EM structure of the (GR)_20_ bound *Oryctolagus cuniculus* ribosome complex PDB:7TOR, https://www.rcsb.org/structure/7TOR, omitting (GR)_20_ and P-tRNA, was used as the starting model for structure refinement for all maps. Domains (60S, 40S head with mRNA, 40S body, E-tRNA, L1 stalk and P stalk) of the 80S were rigid-body fitted into each cryo-EM map using Chimera ^73^. The P-tRNA structure was modeled from PDB: 5UYM, https://www.rcsb.org/structure/5UYM ^57^. Angiogenin was modeled from PDB:5EOP, https://www.rcsb.org/structure/5EOP ^53^, and was remodeled in Coot (vs. 0.8.2) ^74^ to fit the density in the 80S•angiogenin map. Yeast tRNA^Ala^ with the AGC anticodon was modeled in Coot (vs. 0.8.2), using yeast tRNA^Phe^ crystal structure as a starting model (PDB: 1EHZ, https://www.rcsb.org/structure/1EHZ; ^75^). The yeast eEF1A starting model was from AlphaFold (https://alphafold.ebi.ac.uk/entry/P02994, ^76,77^) and the interaction of acceptor arm of tRNA^Ala^ with eEF1A was modeled from the ribosome-bound ternary complex in PDB: 5LZS, https://www.rcsb.org/structure/5LZS ^56^.

The models were refined using real-space simulated-annealing refinement using RSRef ^78,79^ against corresponding maps. Refinement parameters, such as the relative weighting of stereochemical restraints and experimental energy term, were optimized to produce the optimal structure stereochemistry, real-space correlation coefficient and R-factor, which report on the fit of the model to the map ^80^. Harmonic restraints were used for the lower-resolution features of tRNA and eEF1A to maintain secondary structure. The structures were next refined using phenix.real_space_refine ^81^ to optimize protein geometry and B-factors. The resulting structural models have good stereochemical parameters, characterized by low deviation from ideal bond lengths and angles and agree closely with the corresponding maps as indicated by high correlation coefficients and low real-space R factors (**Table S1**). Figures were prepared in Pymol ^82^ and Chimera ^73^.

To visualize the dynamics of tRNA and the P-stalk (**Fig. 3d**), cryo-EM maps from the 6-way classification that yielded the 80S-angiogenin-tRNA^Ala^ were low-pass filtered to 6 Å and rigid-body fitted in Chimera ^73^ using the final 80S-angiogenin-tRNA^Ala^ model (see maps and fits in **Movie S1 and Movie S2)**. For **Fig. 3d**, these rigid fit models were superimposed via the 28S rRNA excluding the flexible P-stalk and L1 stalks and the differences in tRNA positions at the acceptor arm and tRNA elbow were measured in Pymol.

### Ribosome-stimulated nuclease assays

To ascertain whether angiogenin is activated by ribosome binding, we tested tRNA cleavage by angiogenin under conditions similar to those used for assembling the angiogenin bound cryo-EM complexes, using the same preparations of ribosomal subunits, mRNA, angiogenin, and tRNAs, at varied relative concentrations.

To assemble the 80S ribosomal complex, 60S subunits were diluted to 1 µM in 1×10 Buffer (50 mM Tris-Acetate pH 7.0, 100 mM KOAc, 10 mM Mg(OAc)_2_) with 10 mM βME and the mixture was incubated for 5 minutes at 37°C. In a separate microcentrifuge tube, 40S subunits were mixed with mRNA (described in cryo-EM section; Integrated DNA Technologies, Inc, Coralville IA) in the same buffer and incubated for 5 minutes at 37°C then with tRNA^fMet^ for 1 minute, yielding the following concentrations: 1 µM 40S, 20 µM mRNA, and 1.2 µM tRNA^fMet^. The 40S and 60S solutions were mixed 1:1 and the 80S solution was incubated for 5 minutes at 37°C. The solution was diluted to 200 nM 80S for use as a 10x stock in tRNA cleavage assays.

To assemble different ternary complexes, eEF1A, IVT tRNA^Ala^ or Ala-tRNA^Ala^, and GTP or GDPCP were assembled in 1×10 buffer with 10 mM βME by preincubation of eEF1A and GTP (Promega^TM^ Corp., Madison, WI) or GDPCP (Jena Bioscience, Jena, Germany) for 5 minutes at 37°C followed by addition of tRNA^Ala^ or Ala-tRNA^Ala^ for 1 minute at 37°C and were then kept on ice until use. The resulting ternary complex reactions contained 10 µM eEF1A, 2.5 µM tRNA^Ala^ or Ala-tRNA^Ala^, and 1 mM GTP or GDPCP and were used as a 10x stock in tRNA cleavage assays.

The tRNA cleavage reactions were mixed on ice in 1×10 Buffer with 10 mM βME and TritonX and were carried out at 37°C. Order of addition was 1×10 buffer, TritonX (to 0.01% v/v final concentration), rRNasin (to 0.8 U/ul), 80S ribosomal complex (to 20 nM), angiogenin (to 10 nM), then to start the reaction either tRNA^Ala^ or Ala-tRNA^Ala^ or eEF1A•tRNA^Ala^•GTP or eEF1A•Ala-tRNA^Ala^•GTP or eEF1A•Ala-tRNA^Ala^•GDPCP as noted (all tRNA concentration to a final concentration of 250 nM). Upon tRNA addition, the tubes were shifted to 37°C for the indicated times (1, 10 or 100 minutes), whereupon a 10 µl aliquot of the reaction was removed and quenched with 12 µl of 8M Urea Loading Buffer (8M Urea, 2 mM Tris-HCl pH 8, 20 mM EDTA). When all reactions were completed, the reactions were heated at 95°C for 5 minutes, spun down, vortexed and loaded onto a 15% pre-cast Novex Urea-TBE gel (Thermo Fisher Scientific Corp, Waltham MA) and run at 180V for 60 minutes alongside Small RNA Marker (Ab Nova GmbH, Taipei City, Taiwan). The gels were stained in 1xTBE with 1:20,000 SYBR^TM^ Gold nucleic acid stain (Thermo Fisher Scientific Corp, Waltham MA) for 30 minutes. The stained gels were imaged on a ChemiDoc MP system (Bio-Rad Laboratories Inc., Hercules, CA) using the SYBR^TM^ Gold setting. The resulting raw tiff images were analyzed in Fiji ^83^ using the Band/Peak Quantification Tool Marco ^84^ to measure the signal from the tRNA fragments and from the full length tRNA. For each type of tRNA (tRNA^Ala^ vs. Ala-tRNA^Ala^), the ratio of tRNA fragment signal to full length tRNA signal in the no angiogenin control was used to normalize the ratio for other lanes so that an increase in the fragments registers as a fold increase above the starting materials.

---

**Supplementary Figure 1.**
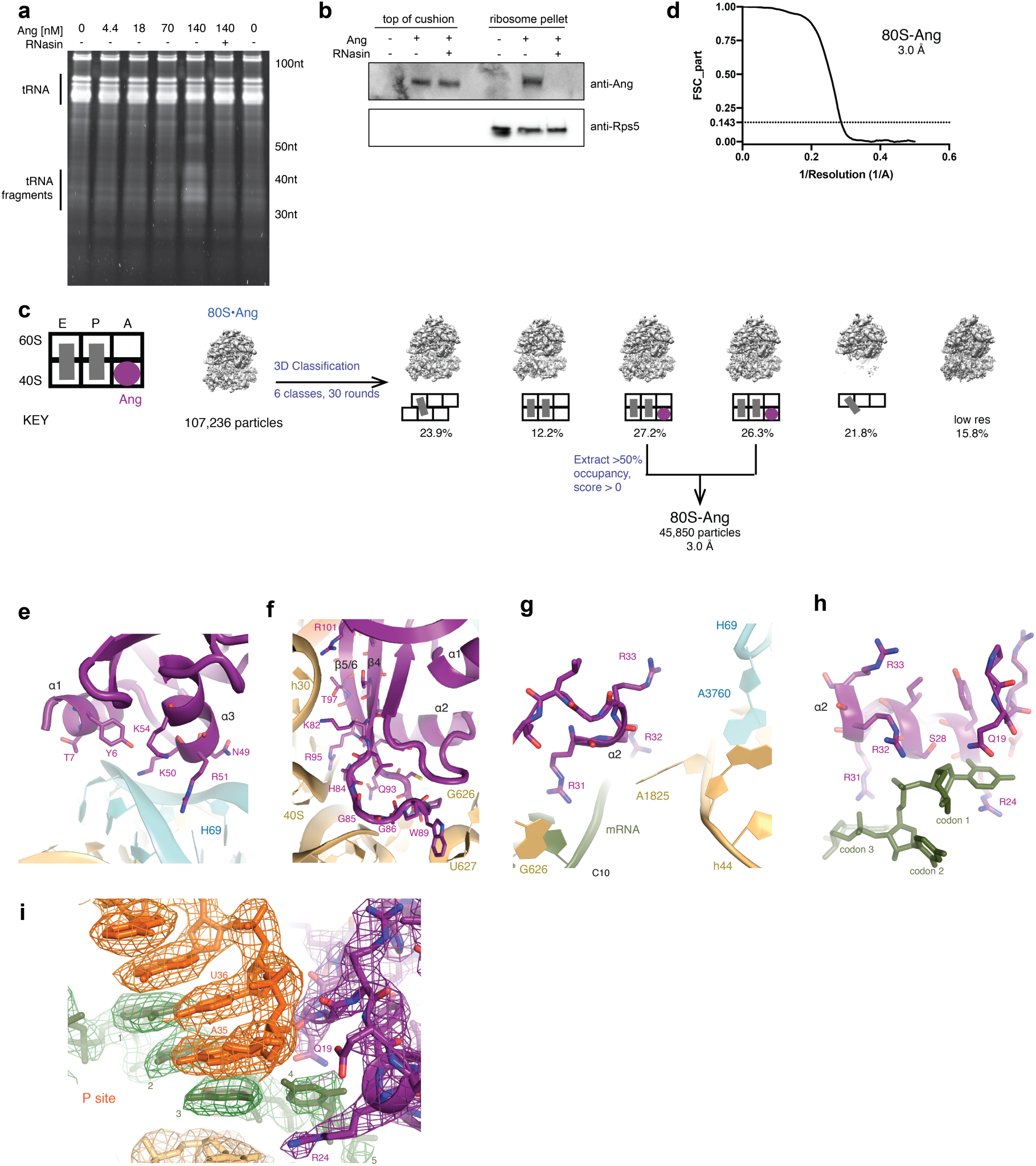
Angiogenin’s activity and binding to the 80S ribosome. (**a**) Incubation of RRL with angiogenin for 15 minutes leads to the accumulation of tRNA fragments; fragment accumulation is inhibited by RNasin; this panel shows an independent repeat of the reactions shown in **Fig. 1b**. (**b**) Angiogenin co-pellets with the ribosome fraction in RRL reactions, except in the presence of RNasin; independent repeat of experiments shown in **Fig. 1c**. (**d**) Maximum-likelihood classification in Frealign of a dataset of 80S ribosome complexes with angiogenin reveals angiogenin bound to the A site of ribosomes in the non-rotated (classical) state. (**d**) Masked, Fourier shell correlation curve as a function of inverse resolution for the map derived from 45,850 particles shown in panel **c** (FSC_part from Frealign v9). (**e**) Interaction of angiogenin with H69. (**f**) Interaction of angiogenin with 18S rRNA. (**g**) Interaction of angiogenin with universally conserved decoding residues. (**h**) Interaction of angiogenin with the A site codon. (**i**) Cryo-EM density showing the interactions of angiogenin with the first nucleotide of the A-site codon adjacent to the P-tRNA anticodon and codon interaction. Note that P-tRNA residue 34 is not shown for clarity. The 2.8 Å cryo-EM map was sharpened by a B-factor of -50 Å^2^ and is shown at 4σ.

**Supplementary Figure 2.**
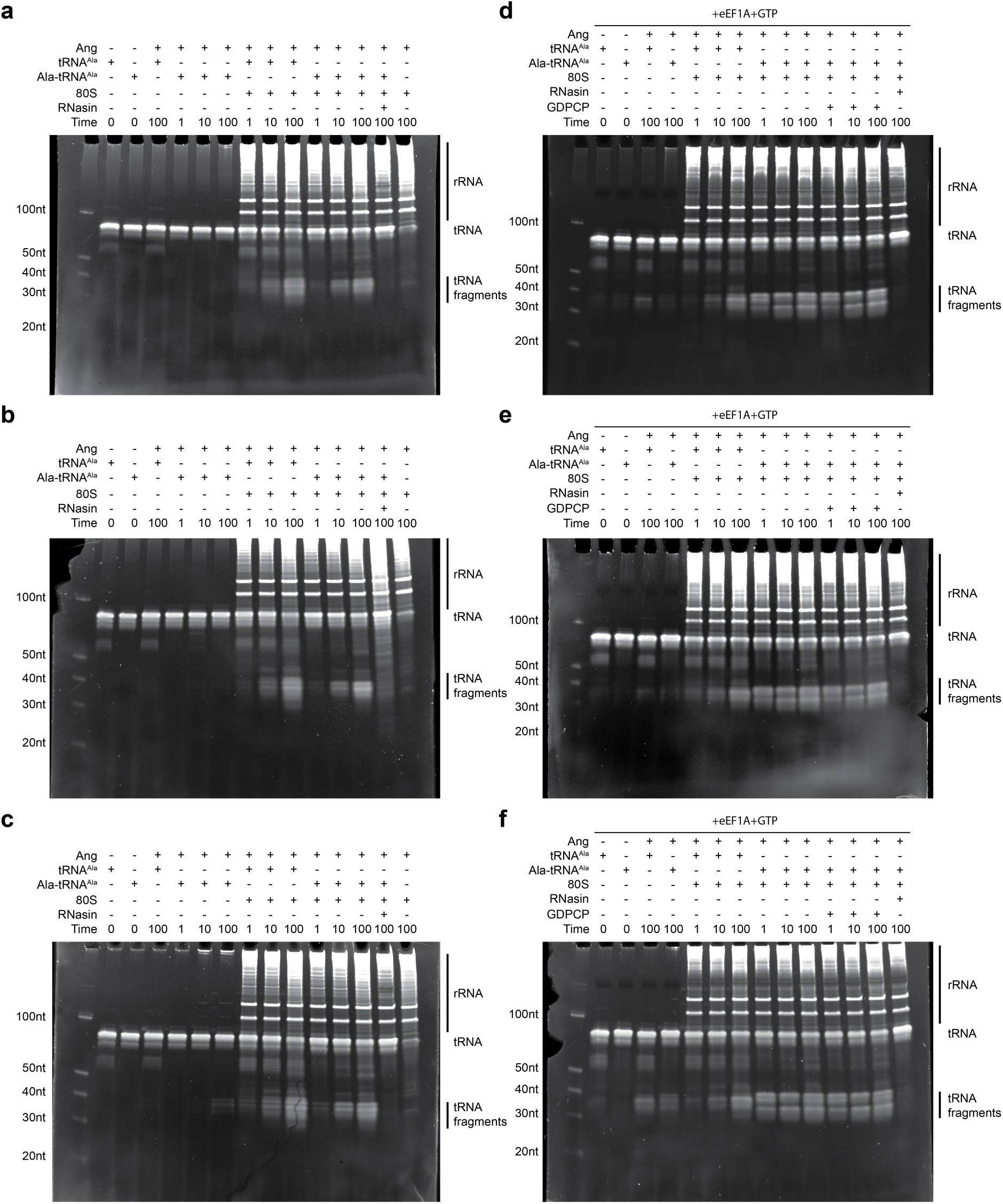
Replicates showing the nucleolytic activity of angiogenin. (**a,b,c**) Production of tRNA fragments by angiogenin is stimulated by the 80S ribosome. 3 independent repeats of tRNA cleavage reactions were separated by denaturing 15% Urea-PAGE gels stained with SybrGold. These gels were quantified for **Fig. 2g**. (**d,e,f**) Production of tRNA fragments by angiogenin is further stimulated by presenting Ala-tRNA^Ala^ as part of a ternary complex with eEF1A. 3 independent repeats of tRNA cleavage reactions were separated by denaturing 15% Urea-PAGE gels stained with SybrGold. These gels were quantified for **Fig. 2i**.

**Supplementary Figure 3.**
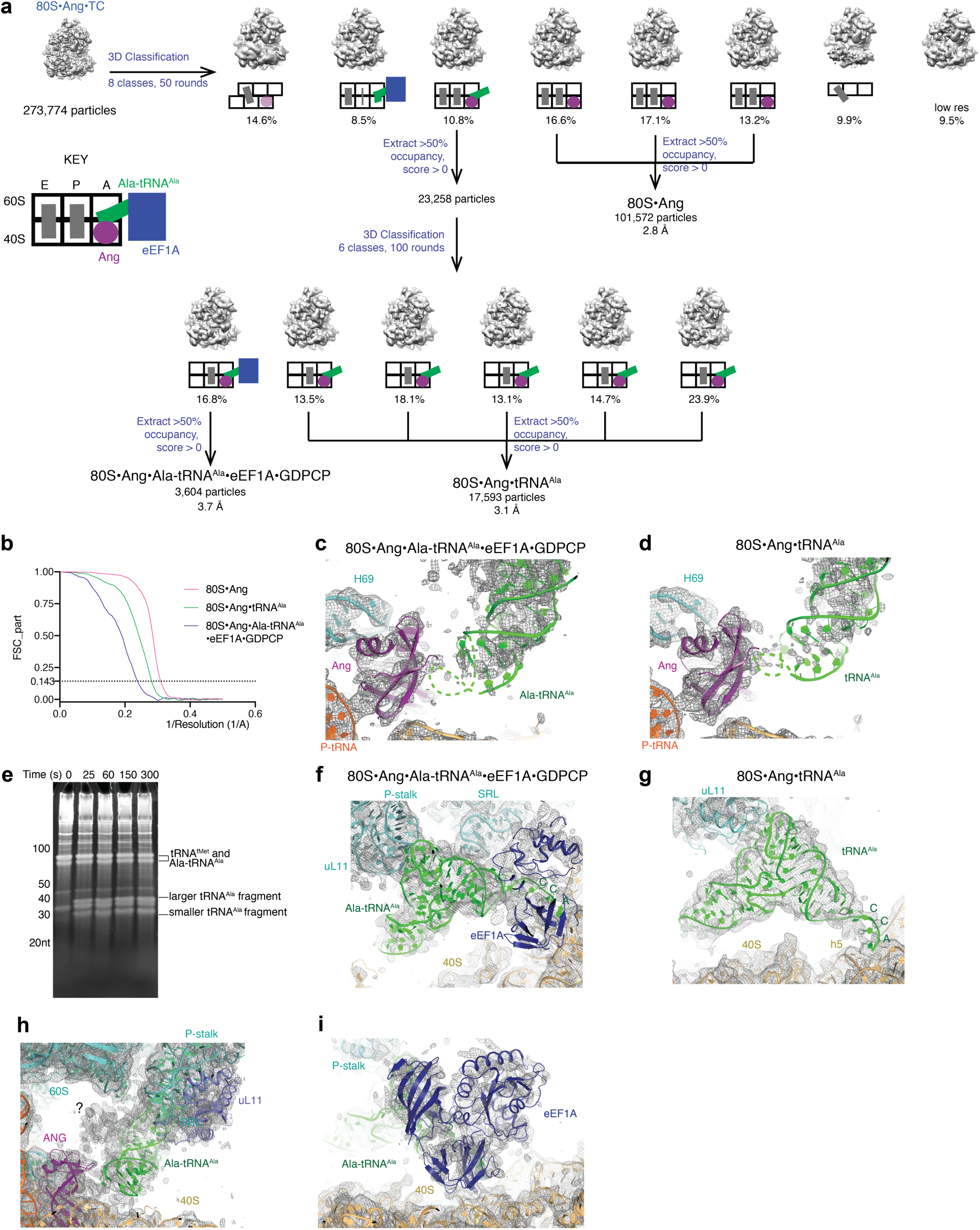
Cryo-EM of the 80S•angiogenin complex with Ala-tRNA^Ala^•eEF1A•GDPCP. (**a**) Maximum-likelihood classification in Frealign of a dataset of 80S ribosome with angiogenin and ternary complex of Ala-tRNA^Ala^•eEF1A•GDPCP shows angiogenin bound to the ribosomal A site as tRNA is presented to the angiogenin active site. (**b**) Masked, Fourier shell correlation curves (FSC) as a function of inverse resolution for the cryo-EM map described in panel **a**. (**c**) The 3.7-Å cryo-EM map of 80S•Ang•Ala-tRNA^Ala^•eEF1A•GDPCP has a well-resolved density for angiogenin and the ribosome but low resolution density for tRNA (see also **Supplementary Fig. 3f**) and nearly absent density for the anticodon loop of Ala-tRNA^Ala^, which was not modeled and is shown as a dashed green line. The cryo-EM map (grey mesh) is shown without sharpening at 4 σ. (**d**) The 3.1-Å cryo-EM map of 80S•Ang•tRNA^Ala^ has well resolved density for angiogenin and the ribosome but low-resolution density for tRNA and weak density for the anticodon loop of tRNA^Ala^ (depicted as a dashed green line). The cryo-EM map (grey mesh) is shown without sharpening at 4 σ. (**e**) Half reactions for assembling the cryo-EM complexes in **Supplementary Fig. 3a** (Half reaction 1: 80S ribosome complexes with angiogenin; Half reaction 2: ternary complex of Ala-tRNA^Ala^•eEF1A•GDPCP as described in Methods) were mixed in test tubes held on wet ice. After the allotted time, the reactions were quenched with 8M Urea loading buffer. For the 0-s time point, Half reaction 1 and Half reaction 2 were added directly to 8M Urea loading buffer. The reactions were separated on a 15% Urea-PAGE gel to visualize full length and newly cleaved tRNA. tRNA is cleaved nearly maximally by 25 s on ice under these conditions, which mimic the conditions used to prepare cryo-EM grids. (**f)** The cryo-EM map of 80S•Ang•Ala-tRNA^Ala^•eEF1A•GDPCP shows an L-shaped density consistent with tRNA interacting with the P-stalk via its elbow and with eEF1A via the CCA end. uL11 and Domain 3 of eEF1A were omitted to show the shape of the tRNA. The cryo-EM map was low-pass filtered to 5 Å and is shown at 3 σ. (**g**) The cryo-EM map of 80S•Ang•tRNA^Ala^ shows an L-shaped density consistent with tRNA interacting with the P-stalk via the tRNA elbow and with h5 of 18S rRNA via the tRNA’s CCA end. uL11 and Domain 3 of eEF1A were omitted to show the shape of the tRNA. The cryo-EM map was low-pass filtered to 5 Å and is shown at 3 σ. (**h**) The cryo-EM map of 80S•Ang•Ala-tRNA^Ala^•eEF1A•GDPCP has an unidentified low-resolution density (question mark) next to the anticodon arm of Ala-tRNA^Ala^ (green) and the P-stalk (uL11, light blue is labeled for reference), which may correspond to a dynamic element of the P-stalk. The cryo-EM map was low-pass filtered to 5 Å and is shown at 2.5 σ. (**i**) Cryo-EM map of 80S•Ang•Ala-tRNA^Ala^•eEF1A•GDPCP shows density for eEF1A (blue) bound to the shoulder of the 40S (gold). The cryo-EM map was low-pass filtered to 5 Å and is shown at 2.7 σ.

**Movie S1**: Cryo-EM maps and rigid-body fits of the subclasses of 80S•Ang•tRNA^Ala^ show different positions of tRNA (green) and P stalk (cyan, upper left corner) relative to angiogenin (magenta). This flexible tethering of the tRNA may allow cleavage at different anticodon positions on various tRNA species. The cryo-EM maps were low-pass filtered to 6 Å and are shown at 2.5 σ.

**Movie S2**: Cryo-EM maps and rigid-body fits of the subclasses of 80S•Ang•tRNA^Ala^ show different positions of tRNA (green) and P stalk (cyan, upper left corner) relative to the 60S subunit’s GTPase activation center. The cryo-EM maps were low-pass filtered to 6 Å and are shown at 2.5 σ.

