## Supplementary figures and images for "Structural mechanism of angiogenin activation by the ribosome"

### Movie S1

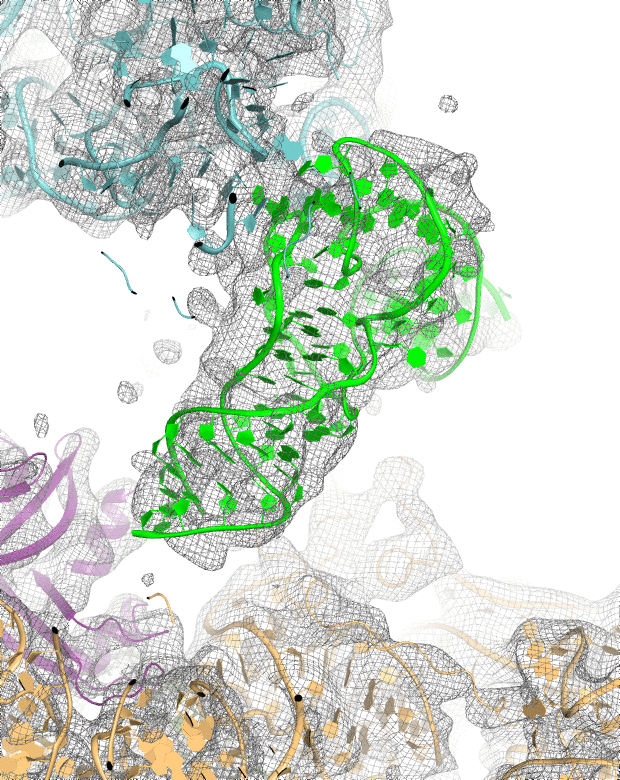

### Movie S2

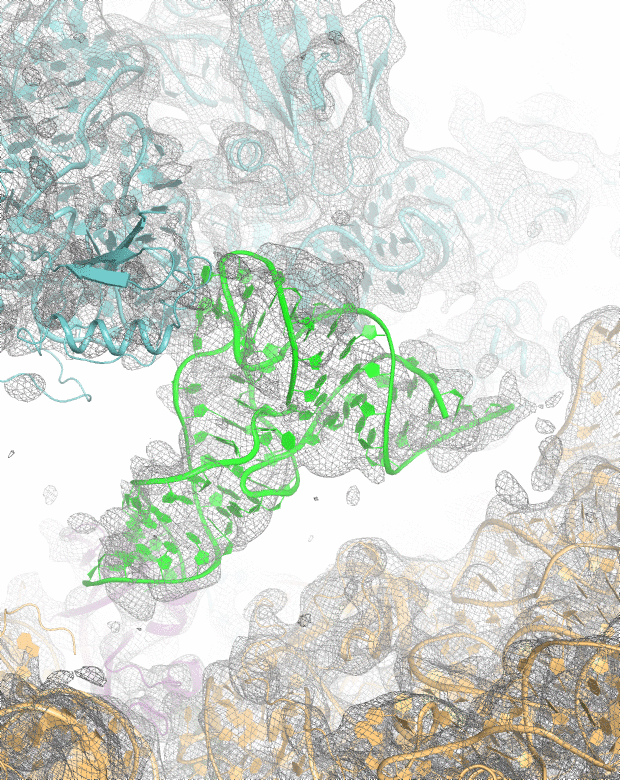
