## Supplementary material for "Structural mechanism of angiogenin activation by the ribosome": Table S1

SUPPLEMENTARY INFORMATION: This file includes:

Table S1

### Table S1: Cryo-EM data collection, refinement and validation statistics

| | 80S Ang alone | 80S Ang | 80S Ang tiRNA | 80S Ang tiRNA eEF1A |
|---|---|---|---|---|
| **Data collection and processing** | | | | |
| Magnification | 57,471x | 57,471x | 57,471x | 57,471x |
| Voltage (kV) | 200 | 200 | 200 | 200 |
| Electron exposure ($e^{-}$/Å$^2$) | 30 | 30 | 30 | 30 |
| Defocus range (μm) | 0.4-3 | 0.4-3 | 0.4-3 | 0.4-3 |
| Pixel size (Å) | 0.87 | 0.87 | 0.87 | 0.83 |
| Symmetry imposed | C1 | C1 | C1 | C1 |
| Initial particle images (no.) | 107,236 | 273,774 | 273,774 | 273,774 |
| Final particle images (no.) | 45,850 | 101,572 | 17,593 | 3,604 |
| Map resolution (Å)** | 3.04 | 2.81 | 3.09 | 3.67 |
| FSC threshold | 0.143 | 0.143 | 0.143 | 0.143 |
| Map resolution range (Å) | 2.7 - >8 | 2.2 - >8 | 2.8 - >8 | 3.0 - >8 |
| **Refinement** | | | | |
| Initial model used (PDB code) | Not modeled | 7TOR | 7TOR | 7TOR |
| Model resolution (Å)* | | 2.8 | 3.1 | 3.7 |
| FSC threshold | | 0.143 | 0.143 | 0.143 |
| Model resolution range (Å) | | 2.4 - >8 | 2.8 - >8 | 3.2 - >8 |
| Correlation Coefficient (cc_mask)* | | 0.90 | 0.87 | 0.80 |
| Real-space R-factor † | | 0.17 | 0.18 | 0.20 |
| Map-sharpening *B* factor (Å$^2$) | | 0 | 0 | 0 |
| Model composition* | | | | |
| Non-hydrogen atoms | | 213,694 | 217,982 | 221,163 |
| Protein residues | | 11,424 | 11,773 | 12,203 |
| RNA residues | | 5,686 | 5,762 | 5,756 |
| *B* factors (Å$^2$)* | | | | |
| Protein | | 114.4 | 129.4 | 152.4 |
| RNA | | 121.1 | 131.4 | 148.2 |
| R.m.s. deviations*,§ | | | | |
| Bond lengths (Å) | | 0.015 | 0.011 | 0.006 |
| Bond angles (°) | | 1.3 | 1.3 | 1.1 |
| Validation* | | | | |
| MolProbity score | | 1.63 | 1.78 | 1.84 |
| Clashscore | | 4.40 | 5.43 | 7.09 |
| Poor rotamers (%) | | 0.0 | 0.02 | 0.01 |
| Ramachandran plot* | | | | |
| Favored (%) | | 93.9 | 91.9 | 93.0 |
| Allowed (%) | | 6.1 | 8.1 | 7.0 |
| Disallowed (%) | | 0.0 | 0.0 | 0.0 |
| Validation (RNA)* | | | | |
| Good sugar pucker (%) | | 98.8 | 99.0 | 99.0 |
| Good backbone (%)# | | 76.2 | 74.2 | 72.9 |

**** from Frealign (FSC_part)**
*** from Phenix**
**† from RSRef**
**§ root mean square deviations**
**# RNA backbone suites that fall into recognized rotamer conformations defined by Molprobity**
